# IFNγ-induced memory in human macrophages is not sustained by epigenetic changes but the durability of the cytokine itself

**DOI:** 10.1101/2025.06.12.659073

**Authors:** Aleksandr Gorin, Noa Harriott, Vyas Koduvayur, Quen J Cheng, Alexander Hoffmann

## Abstract

Macrophages, as key sentinel cells of the innate immune system, can retain memory of prior stimulus exposure. Interferon gamma (IFNγ) plays a central role in maintaining trained immunity *in vivo* and can induce potent memory in macrophages. Such memory is associated with the formation of *de novo* enhancers that alter gene expression responses to subsequent stimuli. However, how such enhancers are maintained after cytokine exposure remains unclear. We report that durable IFNγ-induced enhancers can last for days after cytokine washout, yet the underlying persistence mechanism is not cell-intrinsic. IFNγ-treated macrophages continue to exhibit JAK/STAT signaling days after cytokine removal. Blocking IFNγ signaling with a JAK inhibitor or anti-IFNγ neutralizing antibodies after cytokine removal is sufficient to reverse IFNγ-induced enhancers and erase the potentiated state of the treated macrophages. Our findings suggest that epigenetic changes in macrophages do not inherently encode innate immune memory or a “potentiated” macrophage state, but in fact are themselves dependent on ongoing cytokine signaling. These findings suggest new possibilities for pharmacologic interventions to reverse aberrantly trained immune states associated with pathology.

## Introduction

Innate immune memory, or the ability of the innate immune system to maintain memory of prior immune threats, is apparent in human vaccine cohorts and long-lasting immune sequalae following viral infections(*1–4*). Mice treated with Bacillus Calmette-Guérin (BCG), the fungal compound Beta-D-Glucan (*5–7*) or transient respiratory viral infections (*8–10*) retain improved immunologic responses to subsequent infections even when they lack a functional adaptive immune system. Such memory can be encoded in the bone marrow where hematopoietic progenitor cells differentiate to produce “trained” myeloid cells (*11*). Other models have demonstrated that tissue-resident macrophages at the site of the original exposure are also capable of retaining memory and mediating improved immunologic responses on rechallenge (*8–10, 12*). Recent work has repeatedly demonstrated the central importance of type II interferon, interferon gamma (IFNγ), in such immunologic memory in macrophages (*9, 10, 13–15*).

The mechanism that encodes memory in terminally-differentiated macrophages remains under investigation. However, several *in vivo* studies have demonstrated altered epigenetic landscapes in “trained” tissue-resident macrophages. *In vitro*, macrophages are “polarized” in response to stimuli such as IFNγ; such stimulation of macrophages cells leads to the acquisition of durable enhancer marks (H3K4me1 and H3K4me2) that can persist long after the stimulus is removed (*16–21*). Such *“de novo*” enhancers are thought to mediate long-term memory, mediating potentiated gene expression upon restimulation of the cell (*18, 22, 23*). However, how these histone marks are maintained after stimulus removal remains unknown as the modifications themselves are reversible (*24*).

IFNγ signals by binding to its cognate receptor and activating JAK1/2 signaling which in turn phosphorylate the STAT1 protein. Phosphorylated STAT1 forms homodimers (also known as gamma associate factor, GAF) which translocate to the nucleus and induce the expression of interferon stimulated genes (ISGs), including interferon regulatory factor 1 (IRF1). We have previously demonstrated that IRF1 and GAF work in concert to remodel chromatin and lead to the formation of hundreds of *de novo* enhancers in murine and human macrophages (*18*). We also showed that IFNγ-pulsed macrophages remain hyperresponsive upon restimulation with lipopolysaccharide (LPS) days after the cytokine is removed. Here we report on the mechanism that provides durability to these IFNγ-induced epigenetic changes and resulting capacity for potentiated gene expression responses.

## Results and Discussion

We first asked whether human monocyte-derived macrophages gain enhancer marks in a stimulus-specific manner. After validating the reliability of the Cleavage Under Targets and Tagmentation (CUT&Tag) assay (Supplementary Figure 1), we stimulated human macrophages with IFNγ and LPS for 8 hours and performed H3K4me1 CUT&Tag. LPS activates cells via direct TLR signaling as well as the interferon-beta (IFNβ)-JAK/STAT signaling axis. To identify JAK/STAT-dependent LPS enhancers we treated macrophages (Figure 1A) with LPS in the presence of the JAK inhibitor ruxolitinib at a dose (1µM) sufficient to block IFNβ JAK/STAT signaling (Supplementary Figure 2). We were able to identify 865 IFNγ and 1022 LPS-induced *de novo* enhancers (induced H3K4me1 peaks). Unsupervised *k-means* clustering separated the *de novo* enhancers into 3 major groups (Figure 1B): an LPS-specific/JAK-independent cluster, a cluster of JAK-dependent enhancers shared by LPS and IFNγ, and an IFNγ-specific cluster. Motif analysis of each cluster showed that the most-enriched motif in the LPS-specific/JAK-independent cluster was the NFκB “REL” class while the “IRF1” motif was most highly enriched for JAK-dependent LPS cluster and IFNγ-specific cluster (Figure 1C).

**Figure 1:**
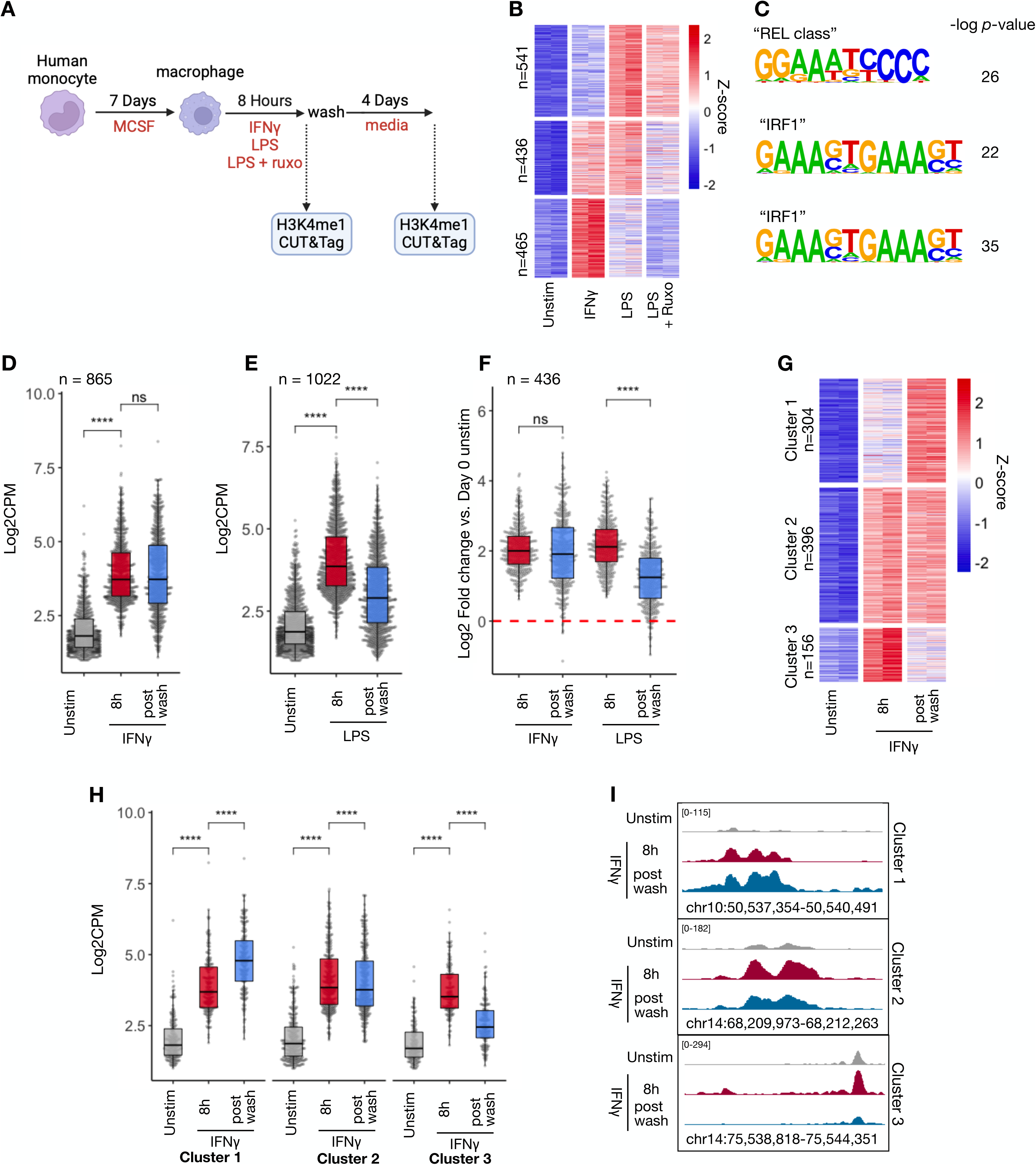
LPS and IFNγ both generate stimulus-specific *de novo* enhancers in human macrophages, however only IFNγ-induced enhancers are durable. A. Schematic of experimental design: Human macrophages were stimulated with either IFNγ (100ng/mL), LPS (100ng/mL), or LPS in the presence of 1µM ruxolitinib for 8 hours. Cells were subsequently washed and cultured for an additional 88 hours. H3K4me1 CUT&Tag was performed at each time point. B. Heatmap of Z-scored reads within H3K4me1 peaks induced by either LPS or IFNγ (L2FC >2, FDR < 0.01). Clusters were generated by unsupervised *k*-means clustering. Each column represents a biological replicate from the same human donor. C. Top enriched motifs in Clusters from (B) D. Box/whisker plot quantifying log2 cpm of reads within IFNγ-induced peaks before and after cytokine washout. E. Box/whisker quantifying log2 cpm of reads within LPS-induced peaks before and after cytokine washout. F. Box/whisker quantifying log2 cpm of reads within peaks induced by both IFNγ and LPS (L2FC >2, FDR <0.01 for each) peaks before and after cytokine washout. G. Z-scored heatmap of reads within CUT&Tag peaks of only IFNy-induced peaks before and after cytokine washout. H. Boxplot of log2cpm of reads within peaks for each cluster identified in (G) I. Examples genome browser tracks for each cluster in E. J. Fraction of peaks containing indicated motif for IFNγ-induced peaks after 8 hours stimulation (L2FC >2, FDR <0.01 vs. unstimulated macrophages) and amongst IFNγ- induced peaks that persist after washout (L2FC ≥0, FDR <0.01). Experiment was performed on two separate human subjects two dots for each motif correspond to each subject. Box/whisker plots indicate interquartile range and 1.5x interquartile range. Statistical tests determined by paired Wilcoxon test. **** *p*<0.0001

Next, we explored enhancer durability after stimulus withdrawal. Macrophages were washed after IFNγ or LPS stimulation and the cells cultured for an additional 88 hours (4 days) in fresh media before performing H3K4me1 CUT&Tag. IFNγ-induced *de novo* enhancers showed persistence after washout (Figure 1D), in contrast, we observed that LPS-induced *de novo* enhancers showed a significant decrease back towards baseline after the stimulus was removed (Figure 1E). We asked whether *de novo* enhancers shared between the two stimuli behave differently after washout; limiting our analysis to the 436 *de novo* enhancers that were induced by both LPS and IFNγ confirmed that IFNγ-induced enhancers persisted (mean L2FC = 0.08, *p* = 0.1) after washout while LPS-induced enhancers reverted towards baseline (mean L2FC=-0.77, *p* < -2^-16)^ (Figure 1F). Further, unsupervised *k*-means clustering of the IFNγ-induced enhancers demonstrated three major patterns of behavior after cytokine washout (Figure 1G-I): one cluster where enhancer marks persisted after washout, one where they decreased after washout, and a third that showed a further increase in the H3K4me1 signal.

To determine whether chromatin opening precedes enhancer formation, as it does in murine BMDMs (*16, 18, 19*), we performed ATACseq on human macrophages stimulated with IFNγ, LPS, or LPS in the presence of ruxolitinib. The results revealed stimulus-specific patterns of chromatin opening, with 7,619 peaks induced by IFNγ and 6,896 by LPS (Figure 2A). Unsupervised *k*-means clustering identified LPS-specific/JAK-independent peaks, peaks shared by LPS and IFNγ that are JAK-dependent, and IFNγ-specific peaks. Motif analysis showed the “NF-kB p65” motif as most enriched in LPS-specific/JAK independent clusters, while “IRF1” was most enriched in IFNγ-specific clusters and JAK-dependent LPS cluster (Figure 2B).

**Figure 2:**
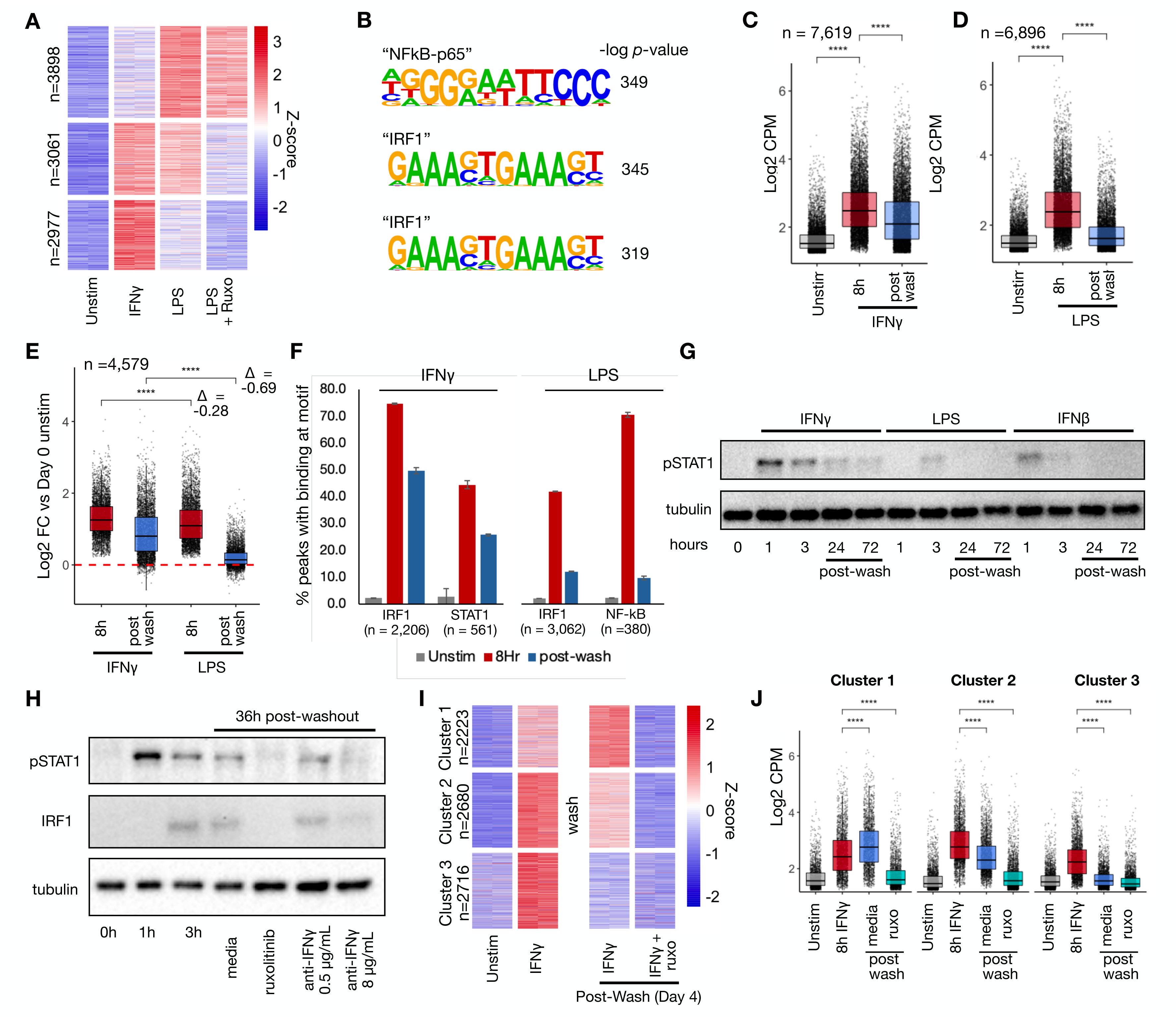
IFNγ induces long-lasting transcription factor activity, and chromatin accessibility after washout, however durability is dependent on continued JAK/STAT signaling. Macrophages were treated with LPS, IFNγ, and LPS in the presence of ruxolitinib for 8 hours as in Figure 1A. Cells were washed and cultured for an additional 88 hours. ATACseq performed after 8 hours of stimulation and 4 days post-washout. A. Heatmap of Z-scored reads within ATAC peaks induced by either LPS or IFNγ (L2FC >2, FDR < 0.01). Clusters were generated by unsupervised *k*-means clustering. Each column represents a biological replicate from the same human donor. B. Top enriched motifs in Clusters from (A) C. Boxplot quantifying log2 cpm of reads within IFNγ-induced ATAC peaks before and after cytokine washout. D. Boxplot quantifying log2 cpm of reads within LPS-induced ATAC peaks before and after cytokine washout. E. Boxplot quantifying log2 cpm of reads within ATAC peaks induced by both IFNγ and LPS (L2FC >2, FDR <0.01 for each) peaks before and after cytokine washout. F. Barplot quantifying percent of transcription factor-bound motifs within STAT1 and IRF1 (IFNγ) and IRF1 and NF-kB (LPS) within induced ATAC peaks in (C) and (D) for unstimulated, IFNγ/LPS-stimulated macrophages, and stimulated macrophages 4 days after washout. Motif binding predicted using TOBIAS ATACseq footprinting analysis. Results are average of two technical replicates from a single subject, error bars display standard deviation. G. Human macrophages were stimulated with IFNγ (100ng/mL), LPS (100ng/mL), or IFNβ (10ng/mL) for 8 hours, washed, and then cultured for an addition 66 hours. Cells were collected and whole cell western blotting for phosphorylated STAT1 was performed at indicated timepoints. Blot is representative of 3 replicates from 2 separate human donors. H. Human macrophages were stimulated with IFNγ (100ng/mL) for 8 hours, washed, and then cultured in regular media or media containing ruxolitinib (1µM) or increasing concentrations of anti-IFNγ neutralizing antibody for an additional 28 hours. Cells were collected and whole cell western blotting for phosphorylated STAT1 and IRF1 was performed at indicated timepoints. Blot is representative of 2 replicates from separate human donors. I. Heatmap of Z-scored reads within ATAC peaks induced by IFNγ (L2FC >2, FDR < 0.01) after 8 hours of stimulation for 4 days post-washout when cultured in regular media or media with 1µM ruxolitinib. Clusters were generated by unsupervised *k*-means clustering. Each column represents a biological replicate from the same human donor. J. Boxplot of log2CPM of reads within each peaks for each cluster in (I). All box/whisker plots indicate interquartile range and 1.5x interquartile range. Statistical tests determined by paired Wilcoxon test. \*\*\**p*<0.001, \*\*\*\**p*<0.0001

To examine whether the chromatin accessibility was similarly transient as reported for LPS-stimulated BMDMs, cells were washed after eight hours of stimulation and cultured for an additional 88 hours when they were again collected for ATACseq. Both IFNγ (Figure 2C) and LPS-induced (Figure 2D) peaks showed a decrease in ATAC accessibility after stimulus washout, however the decrease was less pronounced with IFNγ. Indeed, peaks shared between both stimuli demonstrated more persistence after IFNγ washout (Figure 2E), whereas post-LPS washout most peaks largely had reverted to baseline (L2FC = -0.41 for IFNγ vs -0.98 for LPS).

Next, we asked whether persistent transcription factor activity post-washout may be mediating persistent chromatin accessibility after IFNγ washout. Transcription factor footprinting analysis by TOBIAS(*25*) demonstrated STAT1 and IRF1 binding at accessible chromatin with the majority of induced binding persisting four days after IFNγ washout (66% and 58%, respectively). In contrast, LPS induced IRF1 and NFκB binding was decreased to only 29% and 14% of their maximum levels 4 days after washout (Figure 2F).

Given the durability of chromatin accessibility induced by IFNγ we examined whether IFNγ-induced signaling in the form of phosphorylated STAT1 or IRF1 expression might persist after cytokine washout. We stimulated macrophages with IFNγ, LPS, and IFNβ or eight hours, washed out the stimulus, and cultured the cells for three days. Immunoblotting showed that acute treatment with each stimulus could induce STAT1 phosphorylation. This phosphorylation did not persist after washout of IFNβ and LPS, however, IFNγ-induced STAT1 phosphorylation persisted for three days after the washout (Figure 2G).

Previous studies have shown that IFNγ has inherent affinity for extracellular proteoglycans and phosphatidylserine present on the cell surface of cells from where it may be slowly released to mediate persistent signaling(*26*). To evaluate whether ongoing IFNγ signaling at the cell surface was sufficient to explain the persistent STAT1 phosphorylation in our experiment, we treated macrophages with IFNγ for 8 hours, followed by a wash and then culture in the presence or absence of ruxolitinib, or anti-IFNγ neutralizing antibody for 36 hours. Persistent STAT1 phosphorylation and IRF1 expression were observed post-washout when cells were cultured in media alone, however, both were abrogated by ruxolitinib or high-dose anti-IFNγ antibody (Figure 2H).

To determine if JAK/STAT signaling was essential for sustaining IFNγ-induced chromatin opening, macrophages were treated with IFNγ for eight hours, washed, and subsequently cultured in regular media or media containing ruxolitinib for another 88 hours. Unsupervised *k*-means clustering revealed three patterns of chromatin behavior post-washout when cultured in media alone: increased opening, persistent slight decrease, and abated opening post-washout (Figures 2I-J). Notably, ruxolitinib treatment post-washout reverted chromatin states to baseline, establishing that continued JAK signaling is required for the persistence of chromatin opening following IFNγ stimulation. We observed remarkable similarity between performing this experiment on macrophages from a separate human donor (Supplementary Figure 3A-B).

We asked whether persistent IFNγ-induced transcription factor activity also sustained the expression of ISGs. RNAseq analysis identified 248 IFNγ-induced genes at eight hours of IFNγ treatment (Supplementary Data 1), with 51 of these genes retaining at least 90% and 82 genes maintaining 20-90% of their expression following 88 hours of washout (Figure 3A-B). Ruxolitinib markedly reduced persistent gene expression with no genes maintaining expression above 90%, and only 24 genes retaining expression levels above 20%. To confirm that persistence of gene expression was dependent on IFNγ-signaling rather than non-specific effects of Ruxolitinib we repeated this experiment using anti-IFNγ neutralizing antibody. qPCR of two persistent genes, IRF1 and IDO1, demonstrated that anti-IFNγ antibody also significantly reduced expression of these genes after washout (Figure 3C-D).

**Figure 3:**
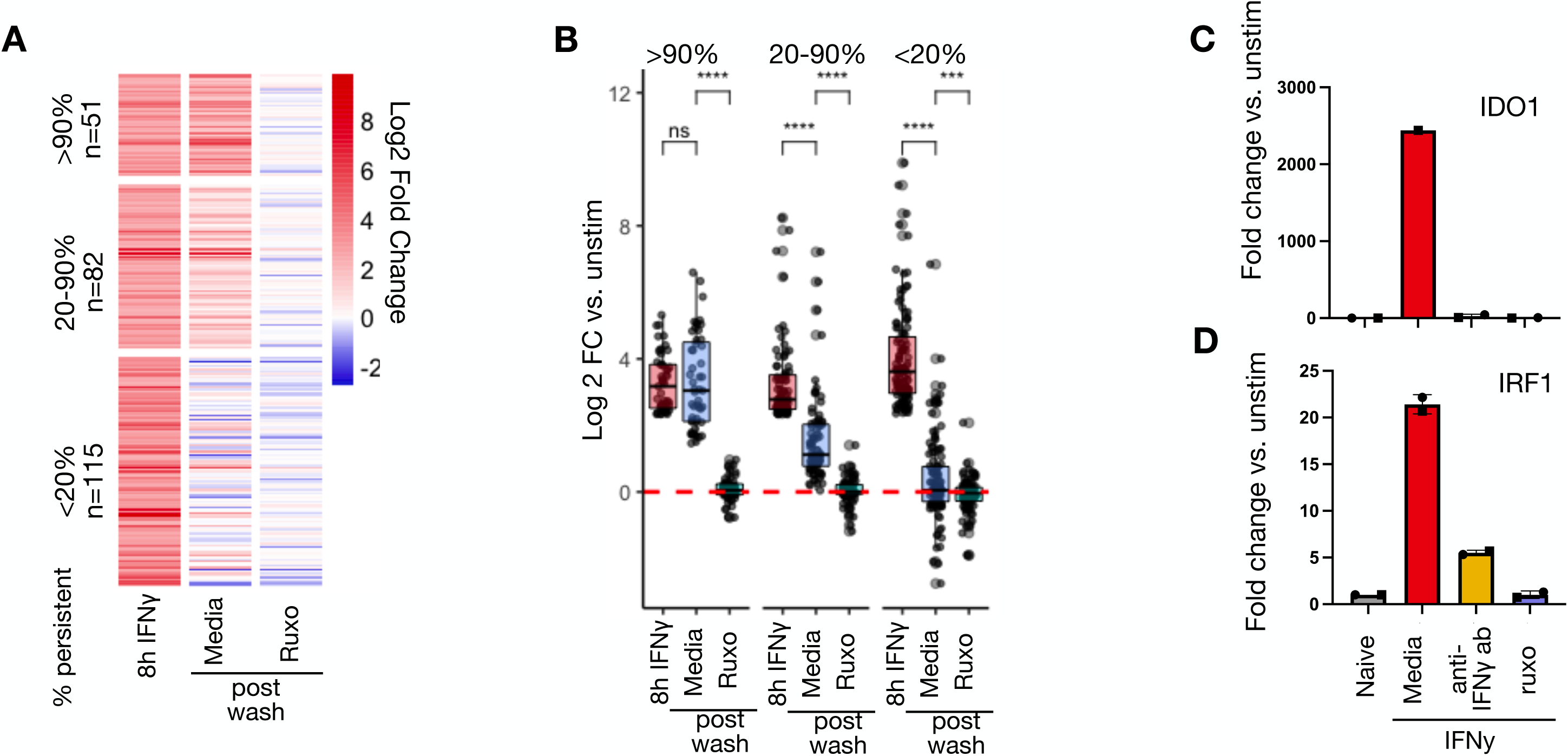
IFNγ signaling sustains ISG expression even after cytokine washout. A. Human macrophages were treated with IFNγ (100ng/mL) for 8 hours, washed, and cultured for an additional 88 hours in the presence and absence of ruxolitinib (1µM). RNAseq was performed after 8 hours of IFNγ-stimulation and 4 days post-washout. Heatmap of Log2 fold change in RNAseq reads of genes induced at least 5-fold after 8 hours of IFNγ stimulation. Log2 fold changes are shown after washout for cells cultured in regular media and media containing 1µM ruxolitinib. Genes are clustered by persistent level of expression post washout (CPM post wash as percent of CPM at 8h simulation). B. Boxplot showing Log2 fold changes of individual genes by cluster in (B). Box/whisker plots indicate interquartile range and 1.5x interquartile range. Statistical tests determined by paired Wilcoxon test. \*\*\**p*<0.001, \*\*\*\**p*<0.0001 C. Macrophages were stimulated and washed as above in (A), after washout cells were cultured in media alone, media with 1µM ruxolitinib, or 8µg/mL anti-IFNγ neutralizing antibody for 88 hours. Cells were collected 88 hours after washout and qPCR performed for IDO1. Boxplots indicate 2^ΔΔCt^ normalized to HPRT. Errors bars indicate standard deviation. D. qPCR for IRF1 as in (C)

We next asked whether persistent signaling is necessary for maintenance of enhancer marks after IFNγ washout (Figure 4A). We found that anti-IFNγ neutralizing antibody and ruxolitinib markedly reduced the persistence of H3K4me1 signals within IFNγ-induced *de novo* enhancers post-washout (Figure 4B). 78.8% of all peaks persisted 88 hours after washout when cells were culture in regular media, compared with 49.6% and 27.5% in media with neutralizing antibody and ruxolitinib, respectively (Figure 4C). Unsupervised *k* means clustering on the H3K4me1 peaks again revealed three patterns of behavior of peaks post-washout: increased reads, unchanged reads, and decreased reads. Addition of neutralizing antibody or ruxolitinib decreased reads in each cluster, demonstrating that the persistence of enhancer marks in each required persistent IFNγ-signaling (Figure 4D-E). We repeated this experiment on macrophages generated from PBMCs from a separate human donor and saw a remarkably similar pattern of enhancer dependence on continued IFNγ signaling (Supplementary Figure 3).

**Figure 4:**
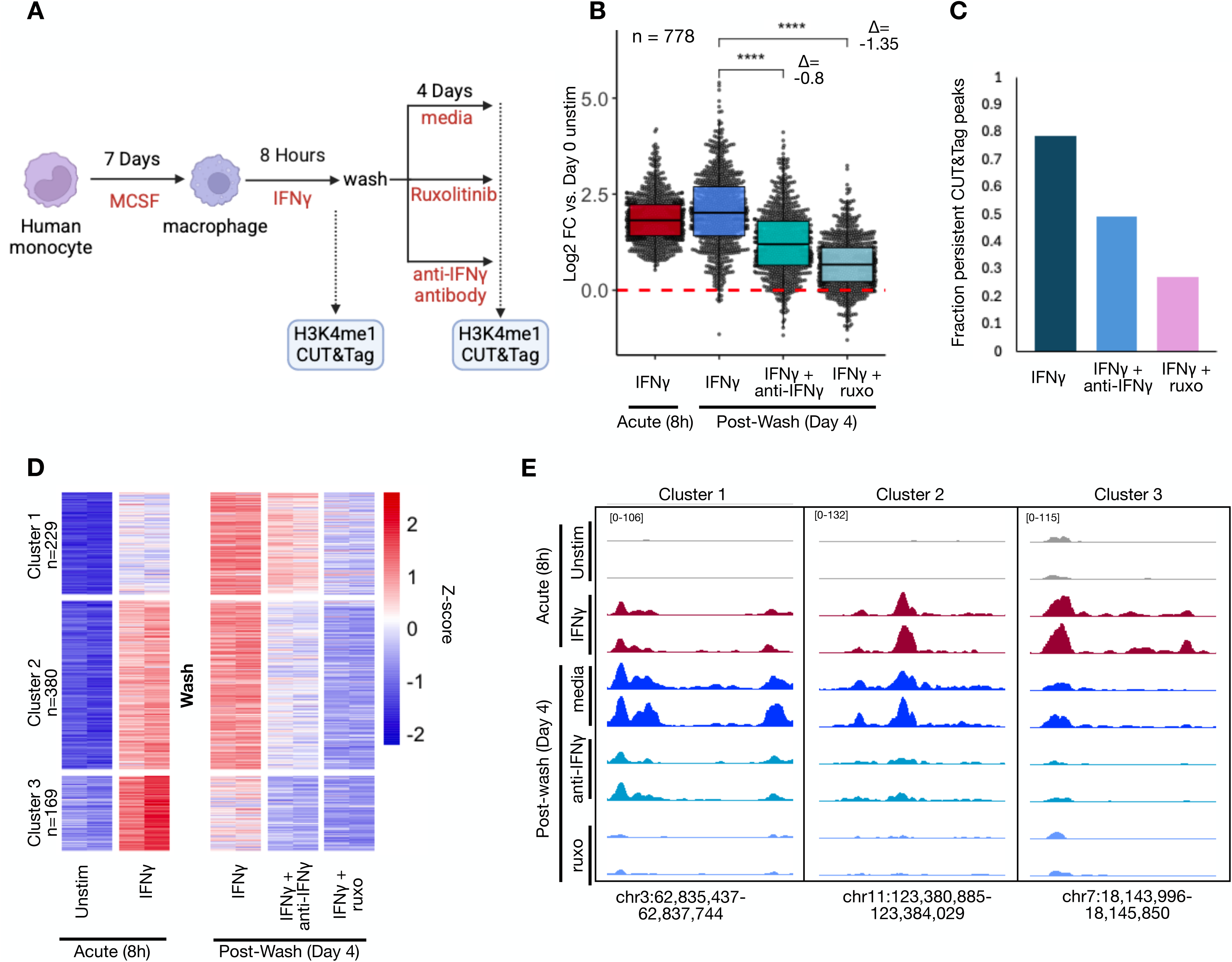
Durability of IFNγ-induced *de novo* enhancers is dependent on continued JAK/STAT signaling by IFNγ. A. Schematic of experimental design: Human macrophages were stimulated with IFNγ (100ng/mL) for 8 hours. Cells were subsequently washed and cultured for an additional 88 hours in standard media, or media supplemented with 1µM ruxolitinib or 8µg/mL anti-IFNγ neutralizing antibody. H3K4me1 CUT&Tag was performed at each time point. B. Boxplot quantifying log2 fold changes of reads within IFNγ-induced H3K4me1 CUT&Tag peaks after 8 hours of IFNγ stimulation and after washout for each condition. C. Barplot showing fraction of IFNγ-induced H3K4me1 peaks at 8 hours that persist 4 days after washout in each condition. Persistence was defined as L2FC ≥0, FDR <0.01. D. Heatmap of Z-scored reads within H3K4me1 peaks induced IFNγ (L2FC >2, FDR < 0.01) after 8 hours of stimulation and 4 days after washout for each condition. Clusters were generated by unsupervised *k*-means clustering. Each column represents a biological replicate from the same human donor. E. Representative genome browser tracks of peaks from each cluster in (D). All box/whisker plots indicate interquartile range and 1.5x interquartile range. Statistical tests determined by paired Wilcoxon test. \*\*\**p*<0.001, \*\*\*\**p*<0.0001

Macrophages pulsed with IFNγ exhibit potentiated expression of inflammatory genes upon exposure to PAMPs such as LPS (*18, 27*). We asked whether sustained signaling was required for maintaining the potentiated state. To this end, we exposed macrophages for eight hrs with IFNγ or vehicle control, cultured them for another four days before exposing them to LPS and collecting samples at one, three, six, and twelve hours of stimulation for RNAseq. We defined potentiated genes as those that displayed at least a five-fold increase in CPM upon LPS stimulation and at least a two-fold greater expression in IFNγ-pretreated cells compared to PBS at two continuous timepoints

Using these criteria we identified 146 LPS-inducible genes potentiated by IFNγ (Supplementary Figure 4B). 40 of these genes had basal expression levels that were equivalent to PBS treated cells (L2FC <0.5 vs PBS treated), while 106 exhibited a higher basal expression after the IFNγ pulse. Potentiated genes with a higher basal set point (e.g. IDO1) exhibited higher expression over the entire LPS time course, while genes with an unchanged basal setpoint (e.g. CSF3) showed potentiation primarily at later timepoints of six and twelve hrs (Supplementary Figure 4 C-E).

Next, we examined if sustained IFNγ signaling is required for the potentiated LPS-response at 4 days after IFNγ washout. To this end we applied ruxolitinib during the washout phase (Figure 5A), but first had to identify LPS-induced genes whose LPS-induction is not blocked by ruxolitinib (Supplementary Figure 2). We identified 45 IFNγ-potentiated LPS-inducible genes whose induction is maintained at least four-fold in the presence of ruxolitinib. Of these, 32 showed elevated basal expression levels (L2FC >0.5) as compared to PBS-treated cells but 13 did not (Figure 5B). Remarkably, ruxolitinib almost entirely abolished potentiated expression, leaving a single gene (*ANKRD1*) still meeting potentiation criteria under JAK blockade (Figure 5C-E).

**Figure 5:**
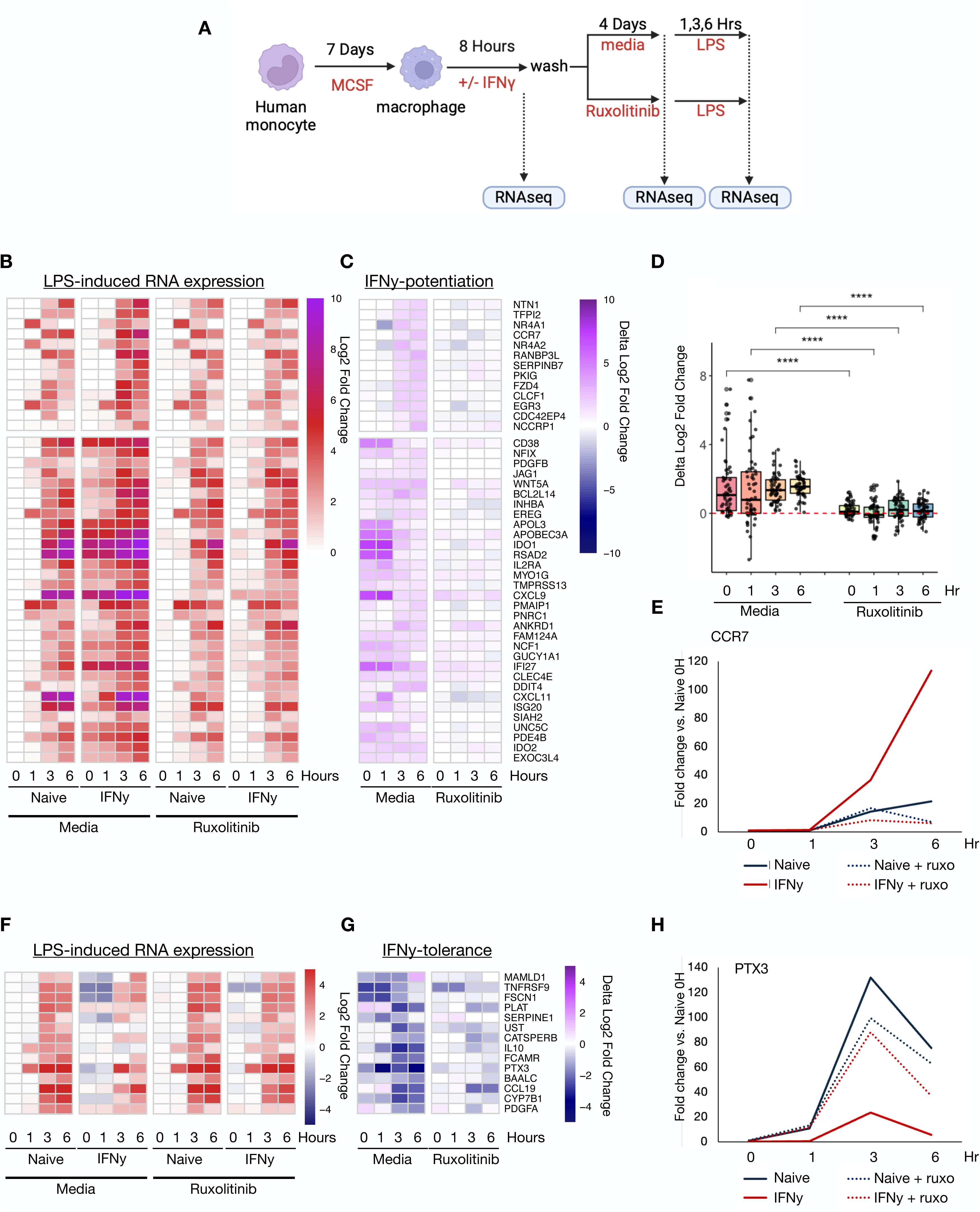
Sustained JAK/STAT signaling is required for long-term IFNγ-induced potentiated and tolerized gene expression responses. A. Human macrophages were stimulated with IFNγ (100ng/mL) for 8 hours. Cells were subsequently washed and cultured for an additional 88 hours in standard media, or media supplemented with 1µM ruxolitinib at which time they were stimulated with 10ng/mL LPS and cultured for an additional 6 hours. RNAseq was performed at each time point. B. Heatmap of log2 fold change in reads of LPS-induced genes potentiated by IFNγ pre-treatment. Log2 fold changes are normalized to PBS-treated controls 88 hours post-washout prior to LPS stimulation (Naïve 0H). Potentiated genes defined as LPS-induced genes reaching at least 4-fold increase for macrophages cultured in ruxolitinib, and at least a 2-fold greater expression in IFNγ-pretreated cells compared to PBS in two contiguous timepoints. Genes are clustered by expression level 88 hours post-IFNγ washout: top cluster of genes showed L2FC <0.5 in IFNγ-treated cells compared to PBS treated, bottom cluster showed >0.5 L2FC those of PBS trained. C. Heatmap quantifying extent of IFNγ-induced potentiation. The difference in L2FC for a given gene between PBS and IFNγ treated is quantified for each gene in (F). D. Boxplot quantifying difference in L2FC for IFNγ pre-treated and PBS-pretreated cells at each timepoint. Box/whisker plots indicate interquartile range and 1.5x interquartile range. Statistical tests determined by paired Wilcoxon test. \*\*\*\**p*<0.0001 E. Example of fold change for potentiated gene: CCR7 F. Heatmap of log2 fold change in reads of LPS-induced genes tolerized by IFNγ. Tolerance is defined as 2-fold reduction in transcription in two contiguous timepoints with IFNγ pre-treatment and at least 4-fold induction by LPS in the presence of ruxolitinib G. Heatmap quantifying extent of IFNγ-induced tolerance as in (C). H. Example of fold change for potentiated gene: PTX3

In addition to potentiating LPS responses, IFNγ may also induce tolerance in some LPS-induced genes (*23*). We examined tolerized genes in our dataset, defined as genes showing a two-fold reduction in transcription in two contiguous timepoints with IFNγ pre-treatment and at least four-fold induction by LPS in the presence of ruxolitinib. This analysis identified 14 genes including well defined tolerized genes IL10 and PTX3 (Figure 5F-E). Ruxotlinib treatment abrogated tolerance of all but two genes (*CCL19, PLAT*).

In sum, the present studies revealed that macrophage memory in response to IFNγ stimulation is not cell-intrinsic, but rather dependent on persistent signaling by the cytokine itself captured on or near the cell surface. In accord with published literature, we demonstrate that IFNγ treatment of macrophages induces GAF/IRF1-driven *de novo* enhancer formation which primes macrophages for heightened responsiveness to LPS stimulation (*18, 22*). However, both the potentiated gene expression capacity and enhancer marks are reversible by inhibiting IFNγ signaling with a neutralizing antibody or a JAK inhibitor, demonstrating that sustained signaling is necessary for their durability.

The persistence of IFNγ signaling days after medium washout extends prior studies that reported that IFNγ has the ability to bind directly to cell membranes and the extracellular matrix (ECM) *via* heparan sulfate and phosphatidyl serine (*26, 28*). These interactions may provide a buffering mechanisms that spatially constrains the cytokine to the site of infection, thereby preventing cytokine overload and systemic IFNγ-induced toxicity (*29*). However, other studies suggested bound IFNγ may be more potent and protected from degradation than soluble cytokine or cytokine in the periphery (*30, 31*). Our work indicates that this spatially constrained IFNγ maintains the IFNγ-polarized epigenetic landscape in macrophages, which in turn maintains potentiated gene expression, thereby facilitating long-term innate immune memory.

Our findings suggest that cytokine-mediated innate immune memory in macrophages is dependent on the surrounding tissue environment rather than being solely intrinsic to the macrophages. This observation is in alignment with recent studies demonstrating that blockade of IFNγ signaling is sufficient to reverse BCG-induced trained immunity in murine models models (*32*). It remains to be seen whether this observation extends to other cytokines or PAMPs/DAMPs that can also induce innate immune memory. Notably, several other cytokines and chemokines have been reported to also have an affinity for the ECM or be spatially constrained *in vivo* (*33–35*). We suggest that acute immune activity within a tissue in response to infection or injury may “stain” the tissue with cytokines and that ongoing signaling from these molecules contributes to lasting changes in tissue resident cells including macrophages. Finally, the observation that the IFNγ-induced memory state is pharmacologically reversible raises the possibility that at least some trained immune states can be pharmacologically erased or modified *in vivo* by blocking cytokine signaling pathways.

## Methods

### Cell culture and stimuli

Human blood from deidentified donors was obtained from the UCLA CFAR Centralized Laboratory Support Core in accordance with UCLA IRB 11-000443. PBMC were isolated from blood by Ficoll (Cytiva 17-1440-03) density centrifugation and cryopreserved in 10%DMSO (Sigma-Aldrich, D2438) in Embryonic Stem cell FBS (Gibco, 11875-093). Monocytes were purified from PBMC using pan-monocyte isolation magnetic beads (Miltenyi 130-096-537). Monocytes were plated on 6-well tissue culture plates at a density of 1.2 x10^6^ cells/well for 6 days in 3mL of RPMI (Gibco, 11875-093) supplemented with 10% ES fetal bovine serum (Gibco, 10439-024), penicillin-streptomycin (Corning, 30-002-CI), L-glutamine (2mM; Corning, 25-005-CI) and human M-CSF (20 ng/ml; R&D systems 216-MC-100). M-CSF was replenished to a concentration of 20ng/mL by adding 60ng M-CSF to each well (assuming that all M-CSF was depleted). On day 7 macrophages were stimulated with human IFNγ (100ng/mL; PeproTech 300-02), LPS (100ng/mL; Sigma-Aldrich, L6529-IMG), human IFNβ1a (10ng/mL; PBL assay science, 11415-1), or PBS as vehicle control for 8 hours. In some conditions cells were pretreated with ruxolitinib (1µM; Selleck S1378) for 15 minutes prior to LPS stimulation. For experiments where cells were washed after stimulation all media was aspirated from the well and each well was thoroughly rinsed three times with 3mL HBSS (Fisher 14-025-134). Following rinsing cells were cultured in complete RPMI media for an additional 36-88 hours as indicated in figures. In some conditions ruxolitinib (1µM) or anti-IFNγ neutralizing antibody (8µg/mL unless otherwise indicated in figure; Invivogen, 10587-44-01) was spiked into complete RPMI media. For RNAseq experiments, macrophages were restimulated 88 hours post-washout with 100ng/mL LPS and samples collected for up to 12 hours post-stimulation.

### CUT&Tag Libraries and Sequencing

Stimulated and control macrophages were lifted from plates with 0.5mM EDTA in PBS and gentle scraping. Nuclear isolation and tagmentation was performed on 100,000 cells per manufacturer protocol as previously described(*36*) with the CUTANA CUT&Tag pAG-Tn5 enzyme (EpiCypher, 15-1017) with the notable exception that a cocktail of 1mM dithiothreitol, 0.5mM (Sigma-Aldrich, D0632), Phenylmethanesulfonyl fluoride (Sigma-Aldrich, 206-350-2), 4µg/mL Leupeptin (Sigma-Aldrich, L9783), 1µM Pepstatin A (Sigma-Aldrich, P5318), and 0.01TIU/mL Aprotinin (Sigma-Aldrich A1153) substituted the Roche Protease inhibitor cocktail where necessary. Anti-H3K4me1 antibody (Abcam, ab8895) was used at a dilution of 1:20 as primary antibody; guinea pig anti-rabbit antibody (Abcam, ABIN101961) was used at a dilution of 1:100 as secondary antibody. Libraries were sequences with paired-end 50bp reads on an Illumina NovaSeq X Plus. Each library was downsampled to 30 Million reads using seqtk with option -s 100. Low quality reads were trimmed (cutoff q=20) and adapter sequences were removed with cutadapt(*37*). Reads were aligned to the human hg38 genome using bowtie2(*38*) with default parameters except –very-sensitive and –non-deterministic options. Aligned reads were filtered based on mapping score (MAPQ ≥30) with Samtools. Duplicated reads were removed with Picard MarkDuplicates. Genome browser tracks were generated using bamCoverage function in deepTools(*39*) with the following options: --binSize 10 --smoothLength 30 --normalizeUsing RPGC --effectiveGenomeSize 2913022398. MACS3(*40*) was used to identify peaks within CUT&tag using standard options except -f BAMPE and -q 0.01. We generated a peak file for every condition and then generated a combined peak file across all conditions for each human subject (including all unstimulated and stimulated conditions). These genomic loci were used to generated a counts table using deeptools multiBamSummary for subsequent analysis. CUT&Tag reads are deposited in the GEO database with accession number GSE294916.

### ATAC Libraries and Sequencing

Stimulated and control macrophages were lifted as described above for CUT&Tag. Nuclear isolation and tagmentation reaction was performed as previously described(*38*). Briefly, 50,000 cells were used to prepare nuclei in cold lysis buffer (10mM Tris-HCl pH 7.5, 3mM MgCl2, 10mM NaCl, and 0.1% IGEPAL CA-630). Nuclei were pelleted by centrifugation at 500g for 10 minutes and suspected in transposase reaction mixture consisting of 25µL 2x TD Buffer, 2.5µL TD enzyme (Illumina, 20034197), and 22.5µL water. The transposase reaction was performed at 37C for 30 minutes in a thermomixer shaker at 800rpm. Libraries were prepared with Nextera DNA library preparation kit and sequences on an Illumina NovaSeq X Plus. Reads were processed and aligned to the human hg38 genome as above for CUT&Tag. ATAC reads are deposited in the GEO database with accession number GSE294915.

### ATACseq and CUT&Tag analysis

Peaks were first filtered to select only those that were in the top 50^th^ percentile of reads in any condition during acute stimulation (unstimulated, 8 hours IFNγ or 8 hours LPS conditions). Differential peaks were identified using edgeR(*41*) applying a cutoff of FDR <0.01 and L2FC >2 compared with unstimulated conditions. Motif analysis to detect top-enriched known motifs was performed with the findMotifsGenome function in the HOMER suite (*42*) using all detected peaks from each condition in each human subject as background. Genome browser tracks were visualized with IGV(*43*) using group auto scale across all conditions for each experiment (ATAC and CUT&Tag were group autoscaled separately).

### TOBIAS transcription factor binding inference

ATAC peaks containing HOMER motifs IRF1 or STAT1 motifs in IFNγ-induced peaks and NFkB-p65-Rel or IRF1 motifs in LPS-induced peaks were identified using the annotatePeaks.pl function from the HOMER suite (v. 4.11). Peaks were filtered and categorized in R by the presence of each motif within the peak within IFNy and LPS induced peaks. TOBIAS (v. 0.17.1) was used to identify regions where transcription factor binding was predicted for each condition. ATACorrect, ScoreBigwig, and BINDetect were run with standard options in sequential order for each condition, input files were bed files of IFNγ and LPS induced peaks and BAM files of aligned reads for each condition. The above homer IRF1, STAT1, and NFkB-p65 motifs were used as input motifs for BINdetect. BINdetect output was used to quantify predicted bound and unbound peaks.

### Immunoblotting

Macrophages were lysed directly on cell culture plates using 2X Laemmli buffer (120mM Tris-Cl, pH = 6.8, 20% glycerol, 4% SDS, 0.05% β-mercaptoethanol, 0.02% bromophenol blue), boiled at 95°C, then stored at -80°C. Equal amounts of protein were loaded in 10% Tris-Glycine gels (Mini-PROTEAN TGX Gels, BioRad #456-1036) separated by molecular weight by electrophoresis at 150 V for 1 hour. Protein was transferred to PVDF membranes at 100 V for 1 hour. Membranes were incubated in 5% bovine serum albumin (BSA, Sigma-Aldrich #A9647) for 1 hour, then incubated in primary antibodies. The following primary antibodies were used: pSTAT1 pY701.4A (Santa Cruz Biotechnology, #136229) diluted at 1:10,000; IRF1 D5E4 (Cell Signaling Technologies, #8478) diluted at 1:1000; β-tubulin TUB2.1 (Sigma-Aldrich, T5201) diluted at 1:10,000. Incubation for pSTAT1 and IRF1 was performed overnight at 4°C, and incubation for β-tubulin was performed for 1 hour at room temperature. Incubation with secondary HRP-conjugated antibodies (anti-mouse IgG, Cell Signaling Technologies #7076; anti-rabbit IgG, Cell Signaling Technologies #7074) was performed for 1 hour at room temperature. Protein was visualized by application of SuperSignal West Pico PLUS Chemiluminescent Substrate (ThermoFisher Scientific #34580) and exposure on a BioRad ChemiDoc MP Imaging System, using BioRad Image Lab software (version 5.2).

### RNAseq sample preparation and analysis

Macrophages were treated as described above. Cells were lysed directly on the plate with Qiagen RLT buffer. RNA was extracted using Qiagen Qiashredder (Qiagen, 79656) and RNEasy mini kit (Qiagen, 74104) according to manufacturer protocol. Library preparation and sequencing was performed by BGI using the DNBseq platform on an MGI T7 machine. Low quality reads were trimmed (cutoff q=20) and adapter sequences were removed with cutadapt. Reads were aligned to the hg38 human genome using STAR(*44*) with the following options: -- outSAMunmapped Within, --outSAMtype BAM SortedByCoordinate, --outFilterType BySJout, -- outFilterMultimapNmax 20, --alignSJoverhangMin 8, --alignSJDBoverhangMin 1, -- outFilterMismatchNmax 999, --alignIntronMin 20, --alignIntronMax 1000000, -- alignMatesGapMax 1000000, --outFilterMismatchNoverLmax 0.04, --seedSearchStartLmax 30. Aligned reads were filtered based on mapping score (MAPQ ≥ 30) by Samtools. Counts for each gene were generated using featureCounts(*45*). Counts were normalized by CPM. Analysis of gene expression was limited to protein coding genes. RNAseq reads are deposited in the GEO database under accession number GSE294918.

### Gene Expression Analysis by RT-qPCR

Macrophages were lysed directly on tissue culture plates and RNA was collected and purified using RNeasy Mini Kit (Qiagen, #74106). Equal amounts of RNA were reverse transcribed into cDNA using LunaScript RT SuperMix Kit (New England Biolabs #E3010). Equal amounts (10 ng) of cDNA were loaded for qPCR analysis, using the Luna Universal qPCR Master Mix (New England Biolabs #M3003) and 0.25 μM each of forward and reverse primers for target genes: IDO1 (5’-GCTGGGACATCAACAAGGAT-3’; 5’-CTTCCACGTCTTGGGATCTG-3’); IRF1 (5’-GCTGGGACATCAACAAGGAT-3’; 5’-CTTCCACGTCTTGGGATCTG-3’); and HPRT1 (5’-AGGACTGAACGTCTTGCTCG-3’; 5’-ATCCAACACTTCGTGGGGTC-3’).

