## Supplementary Figures for "IFNγ-induced memory in human macrophages is not sustained by epigenetic changes but the durability of the cytokine itself"

**Supplementary Figure 1: CUT&Tag identifies H3K4me1 peaks and is distinct from ATAC**
To validate that the CUT&Tag assay for H3K4me1 identifies true histone marks over potential background Tn5 activity, we performed the Assay for Transposase-Accessible Chromatin (ATACseq) on paired samples from the same subject collected simultaneously as the CUT&Tag assay. Peaks were identified based on CUT&Tag reads and reads within the peaks was quantified for both CUT&Tag and ATAC experiments. The results show greater consistency within each assay group rather than across, regardless of treatment). Reads are normalized across ATAC and CUT&Tag independently.

A. Spearman correlation of reads within the same peaks for CUT&Tag and ATAC experiments.
B. Genome browser track of reads within an identified CUT&Tag peak showing minimal reads within the same peak in an ATAC experiment.
C. Genome browser track of reads within an identified CUT&Tag peak showing higher reads within the same peak in an ATAC experiment.
D. Genome browser track of reads within an identified CUT&Tag peak showing similar reads within a peak between ATAC and CUT&Tag experiments.

**Supplementary Figure 2: Ruxolitinib blocks LPS and IFNγ-induced Janus Kinase signaling.**

Human macrophages were pre-treated with increasing concentrations of ruxolitinib for 15 minutes and subsequently stimulated with IFNγ (100ng/mL) or LPS (100ng/mL) for 3 hours. Whole cell lysate western blots showing effect of ruxolitinib on STAT1 and STAT2 phosphorylation by each stimulus. Blot is representative of 2 replicates from separate human donors.

**Supplementary Figure 3: Macrophages from a second human donor show reversibility of IFNγ-induced chromatin accessibility and *de novo* enhancers**
Human macrophages from a second human subject were stimulated with IFNγ (100ng/mL) for 8 hours. Cells were subsequently washed and cultured for an additional 88 hours in standard media, or media supplemented with 1μM. ATAC and H3K4me1 CUT&Tag was performed after 8 hours of stimulations and 88 hours after cytokine washout.

A. Boxplot quantifying log2 fold changes of reads within IFNγ-induced ATAC peaks after 8 hours of IFNγ stimulation and after washout for each condition (L2FC >2, FDR < 0.01).
B. Heatmap of Z-scored reads within ATAC peaks induced IFNγ after 8 hours of stimulation and 4 days after washout for each condition. Clusters were generated by unsupervised *k*-means clustering. Each column represents a biological replicate from the same human donor.
C. Boxplot quantifying log2 fold changes of reads within IFNγ-induced H3K4me1 CUT&Tag peaks after 8 hours of IFNγ stimulation and after washout for each condition (L2FC >2, FDR < 0.01).
D. Barplot showing fraction of IFNγ-induced H3K4me1 peaks at 8 hours that persist 4 days after washout in each condition. Persistence was defined as L2FC ≥0, FDR <0.01.
E. Heatmap of Z-scored reads within H3K4me1 peaks induced IFNγ after 8 hours of stimulation and 4 days after washout for each condition. Clusters were generated by unsupervised *k*-means clustering. Each column represents a biological replicate from the same human donor.
F. Boxplot of log2CPM of reads within each peaks for each cluster in (E).

Box/whisker plots indicate interquartile range and 1.5x interquartile range. Statistical tests determined by paired Wilcoxon test. ****$p$<0.0001

**Supplementary Figure 4: IFNγ exposed macrophages exhibit potentiated inflammatory gene expression upon LPS restimulation**

A. Human macrophages were stimulated with IFNγ (100ng/mL) for 8 hours. Cells were subsequently washed and cultured for an additional 88 hours in standard media, or media supplemented with 1μM ruxolitinib at which time they were stimulated with 10ng/mL LPS and cultured for an additional 12 hours. RNAseq was performed at each time point.
B. Heatmap of log2 fold change in reads of LPS genes potentiated by IFNγ pre-treatment. Log2 fold changes are normalized to PBS-treated controls 88 hours post-washout prior to LPS stimulation (Naïve 0H). Potentiated genes defined as those reaching 5-fold increase in reads post-LPS stimulation and at least a 2-fold greater expression in IFNγ-pretreated cells compared to PBS. Genes are clustered by expression level 88 hours post-IFNγ washout: Top cluster of genes showed L2FC <0.5 in IFNγ-treated cells compared to PBS treated, bottom cluster showed >0.5 L2FC those of PBS trained.

C. Heatmap quantifying extent of IFNγ-induced potentiation. The difference in L2FC for a given genes between PBS and IFNγ treated is quantified for each gene in (F).
D. Example of CPM for potentiated gene that showed basal expression equivalent to that of PBS treated cells: CSF3
E. Example of CPM for potentiated gene that showed basal expression higher than of PBS treated cells: IDO1

### Figure S1

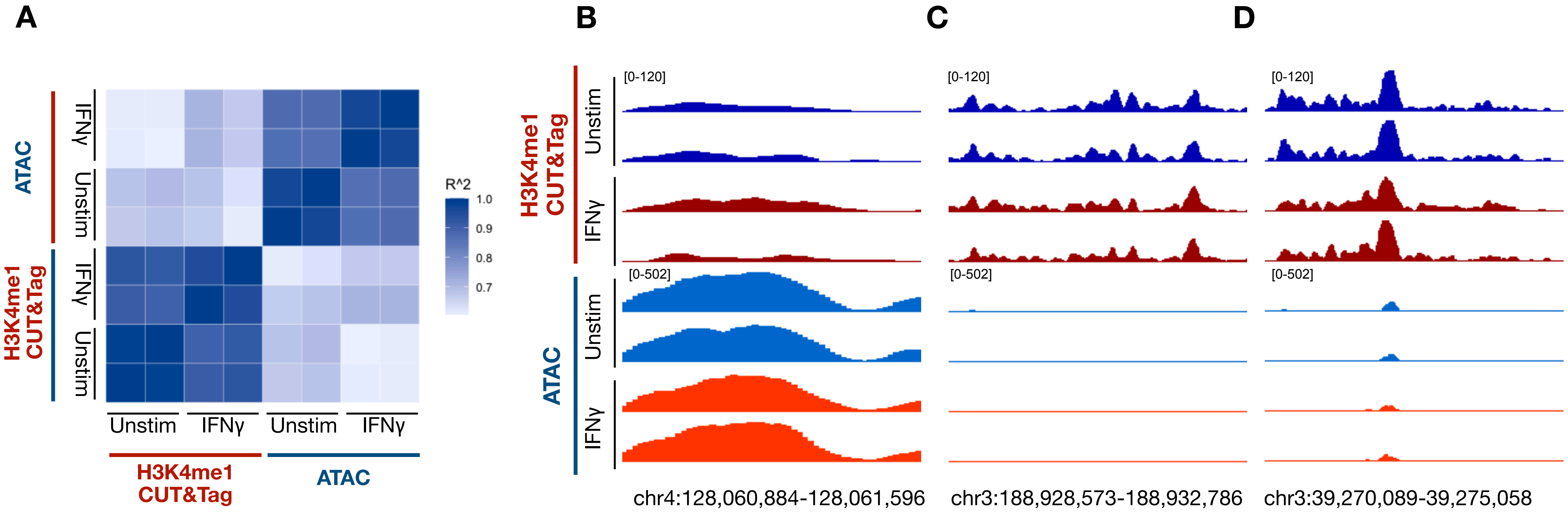

A
B
C
D
ATAC
H3K4me1
CUT&Tag
IFNγ
Unstim
R^2
1.0
0.9
0.8
0.7
[0-120]
[0-502]
chr4:128,060,884-128,061,596
chr3:188,928,573-188,932,786
chr3:39,270,089-39,275,058

**Figure S2**

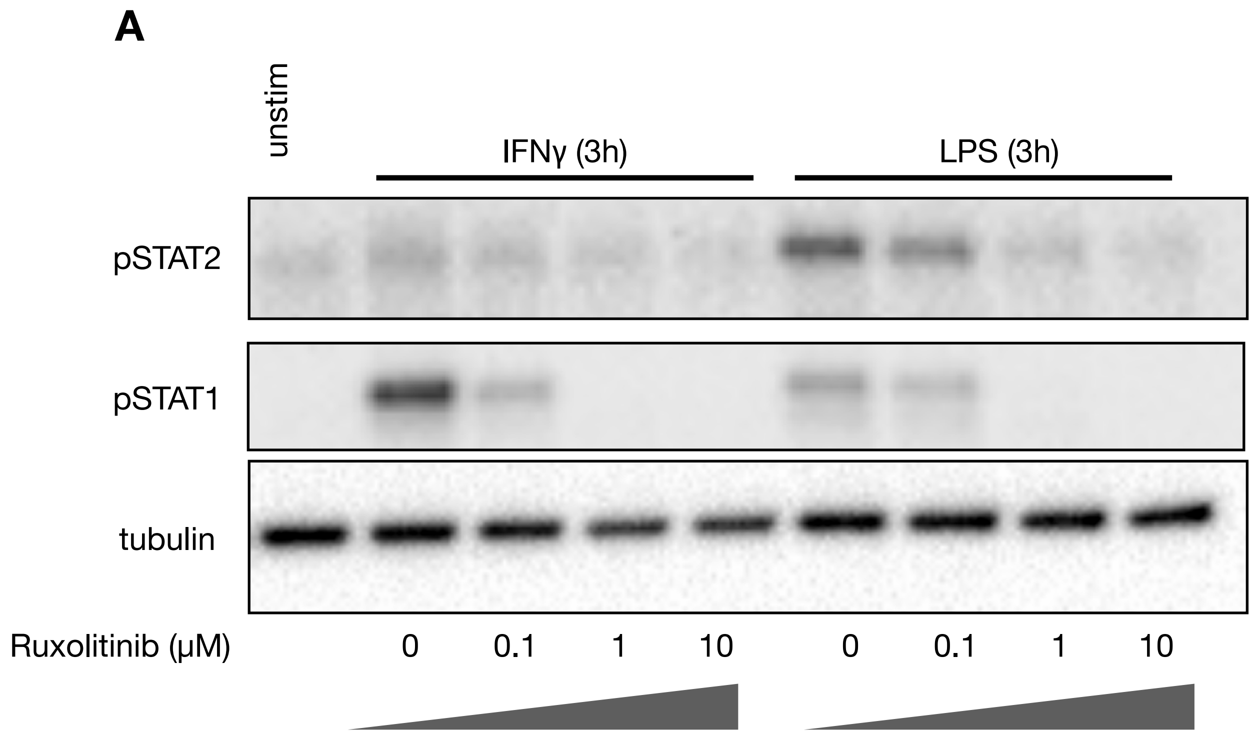
A
unstim
IFNγ (3h)
LPS (3h)
pSTAT2
pSTAT1
tubulin
Ruxolitinib (μM)
0
0.1
1
10
0
0.1
1
10

**Figure S3**

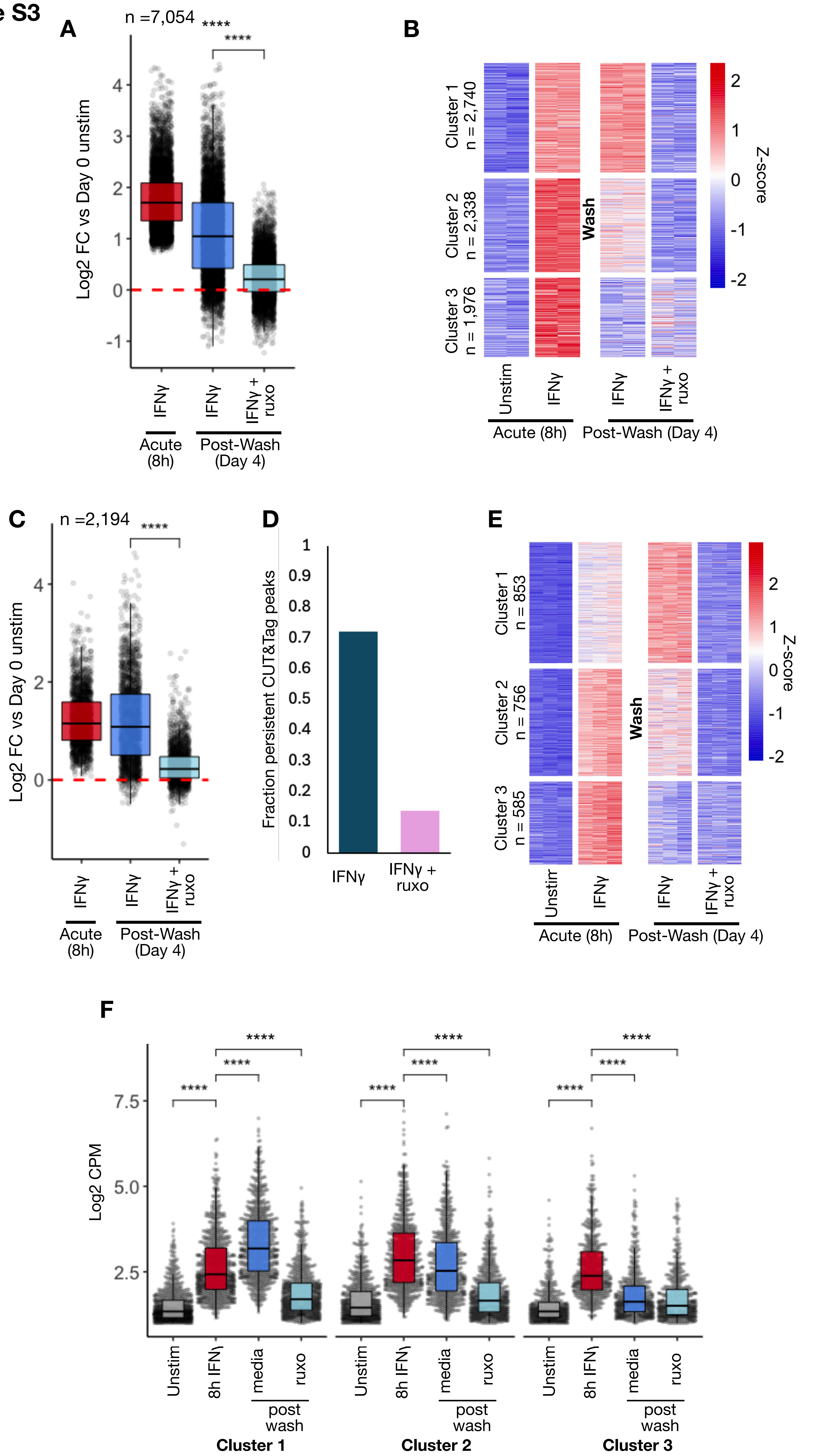

A
n =7,054
****
****
Log2 FC vs Day 0 unstim
IFNγ
IFNγ
IFNγ + ruxo
Acute (8h)
Post-Wash (Day 4)
B
Cluster 1 n = 2,740
Cluster 2 n = 2,338
Cluster 3 n = 1,976
Wash
Z-score
Unstim
IFNγ
IFNγ
IFNγ + ruxo
Acute (8h)
Post-Wash (Day 4)
C
n =2,194
****
Log2 FC vs Day 0 unstim
IFNγ
IFNγ
IFNγ + ruxo
Acute (8h)
Post-Wash (Day 4)
D
Fraction persistent CUT&Tag peaks
IFNγ
IFNγ + ruxo
E
Cluster 1 n = 853
Cluster 2 n = 756
Cluster 3 n = 585
Wash
Z-score
Unstim
IFNγ
IFNγ
IFNγ + ruxo
Acute (8h)
Post-Wash (Day 4)
F
Log2 CPM
****
Unstim
8h IFNγ
media
ruxo
post wash
Cluster 1
Cluster 2
Cluster 3

**Figure S4**

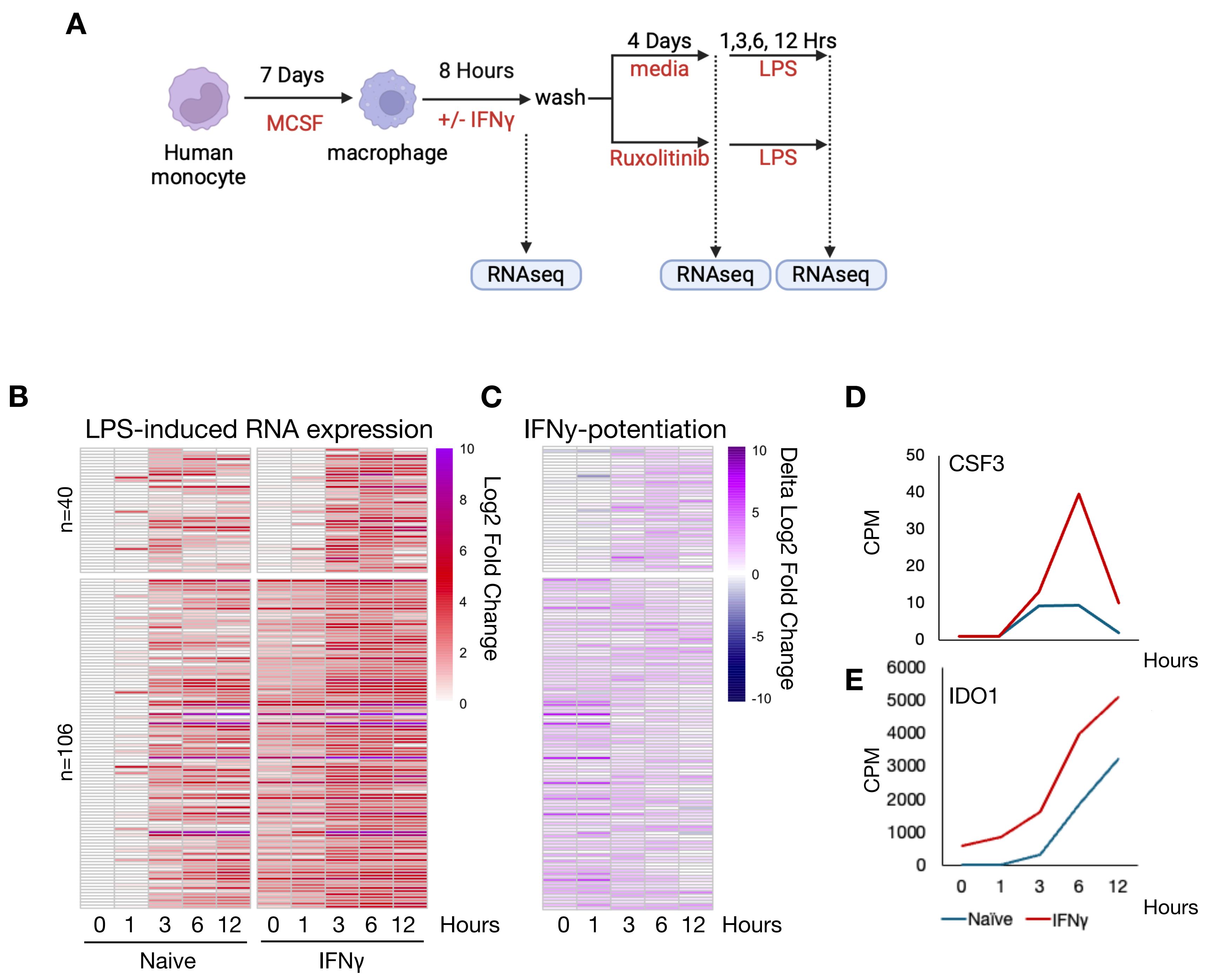

A
7 Days
MCSF
Human
monocyte
macrophage
8 Hours
+/- IFNγ
wash
4 Days
media
Ruxolitinib
1,3,6, 12 Hrs
LPS
LPS
RNAseq
RNAseq
RNAseq
B
LPS-induced RNA expression
n=40
n=106
0 1 3 6 12 0 1 3 6 12 Hours
Naive
IFNγ
Log2 Fold Change
10 8 6 4 2 0
C
IFNγ-potentiation
0 1 3 6 12 Hours
Delta Log2 Fold Change
10 5 0 -5 -10
D
CSF3
CPM
50 40 30 20 10 0
Hours
E
IDO1
CPM
6000 5000 4000 3000 2000 1000 0
0 1 3 6 12
Hours
Naïve
IFNγ
